# Methane, a gut bacteria-produced gas, does not affect arterial blood pressure in normotensive anaesthetized rats

**DOI:** 10.1101/2021.03.31.437828

**Authors:** Ewelina Zaorska, Marta Gawryś-Kopczyńska, Ryszard Ostaszewski, Dominik Koszelewski, Marcin Ufnal

## Abstract

Methane is produced by carbohydrate fermentation in the gastrointestinal tract through the metabolism of methanogenic microbiota. Several lines of evidence suggest that methane exerts anti-inflammatory, anti-apoptotic and anti-oxidative effects. The effect of methane on cardiovascular system is obscure. The objective of the present study was to evaluate the hemodynamic response to methane. A vehicle or methane-rich saline were administered intravenously or intraperitoneally in normotensive anaesthetized rats. We have found no significant effect of the acute administration of methane-rich saline on arterial blood pressure and heart rate in anaesthetized rats. Our study suggests that methane does not influence the control of arterial blood pressure. However, further chronic studies may be needed to fully understand hemodynamic effects of the gas.

## Introduction

Methane (CH_4_) is the simplest alkane and the most abundant organic gas in the atmosphere. It is a combustible, intrinsically non-toxic gas. While methane is intrinsically non-toxic, it is extremely flammable. Mixing methane with atmospheric air results in an exothermic reaction. As such, methane’s easily combustible nature and reactivity may lead to explosions. However, when controlled, these properties make methane an excellent source of fuel. It also plays a role in global warming. [1,2]

CH_4_ can be produced by carbohydrate fermentation in the gastrointestinal (GI) tract through the metabolism of methanogenic microorganisms. The catalyzing enzyme of this pathway is methyl coenzyme M reductase. [3,4] Mammalian methanogenesis is not the only source of endogenous CH_4_. Numerous in vitro and in vivo studies have revealed the possibility of non-microbial CH_4_ formation in mitochondria and eukaryotic cells. It was proposed that CH_4_ liberation is related to hypoxic events resulting in, or associated with, mitochondrial dysfunction. [5–9]

In humans, the respiratory system excretes the bulk of the endogenous methane. [10] The intraluminally generated CH_4_ enters the splanchnic circulation and is then released in the breath if the partial pressure is higher than in the atmosphere. [11] Considering the physicochemical properties and the favorable distribution properties of methane, it is assumed that it can also be excreted through other body surfaces. [12]

Due to methane’s flammable and explosive nature, its use by inhalation route in research is limited. [13,14] A more convenient and safer method of delivering methane is to use methane-rich saline (MRS). [15–17]

In recent years, many researchers have focused on methane’s biological activity, especially its anti-inflammatory, antioxidant, and anti-apoptotic properties. [18–21] It turns out that methane produced in the gastrointestinal tract can slow down intestinal transit, augment small intestinal contractile activity and exert a protective effect in ischemia/reperfusion (IR)-induced intestinal injury.[22,23]

It was also found that methane plays a protective role in hepatitis, acute lung injury, myocardial ischemia injury, sepsis and diabetic retinopathy. [24–27] According to other research, methane shows a neuroprotective effect in the rat model of acute carbon monoxide poisoning and spinal cord ischemia/reperfusion injury. [28,29]

Due to its properties, availability and protective effect in many animal models of diseases, it was proposed that methane, similarly to hydrogen sulfide and nitrogen oxide, can act as a gaseous transmitter. Although methane has been studied in many medical fields, little attention has been paid to its role in blood pressure regulation. [30–34]

The objective of the present study was to evaluate the hemodynamic response to methane in normotensive rats.

## Results and Discussion

The mean arterial blood pressure (MABP) and heart rate (HR) were measured at baseline and after the administration of either a vehicle or methane-rich saline (MRS). The methane-rich saline solution was prepared in accordance with the procedure described by Ye et al. [35]

There was no significant difference in the MABP or HR at baseline between the experimental series.

In anesthetized rats, intravenous and intraperitoneal administration of both the vehicle and MRS produced transient and not significant changes in MABP and HR (Figures 1 and 2).

**Fig. 1.**
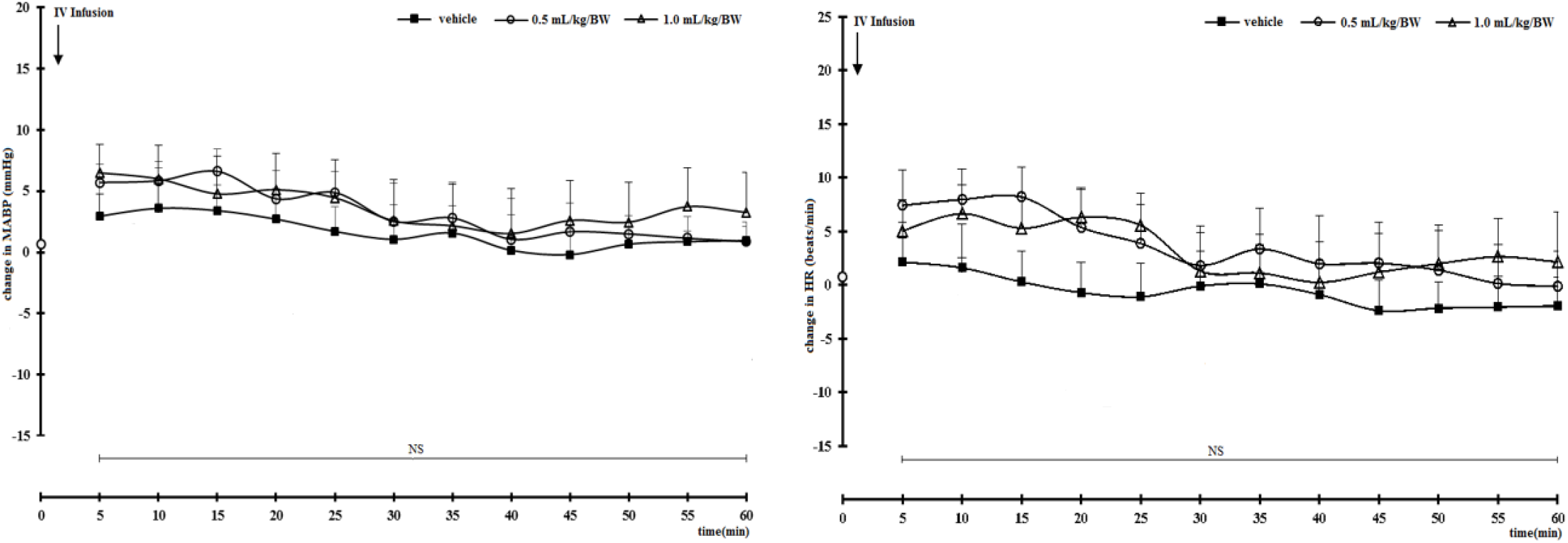
Changes in the mean arterial blood pressure (MABP, mmHg) and heart rate (HR, beats/min) after the intravenous administration of vehicle (saline/ aqueous 0.9 % NaCl n = 5) or methane-rich saline (MRS) [(n = 5) at a dose of 0.5 mL/kg/BW, (n = 5) at a dose of 1.0 mL/kg/BW] in normotensive Sprague Dawley rats. NS (p > 0.05) vs. baseline, NS (p > 0.05) – 0.5, 1.0 mL/kg/BW MS series vs. vehicle, NS (p > 0.05) – 1a (0.5 mL/kg/BW) vs. 1b (1.0 mL/kg/BW) series. NS – not significant, comparisons between the treatments by ANOVA for repeated measures.

**Fig. 2.**
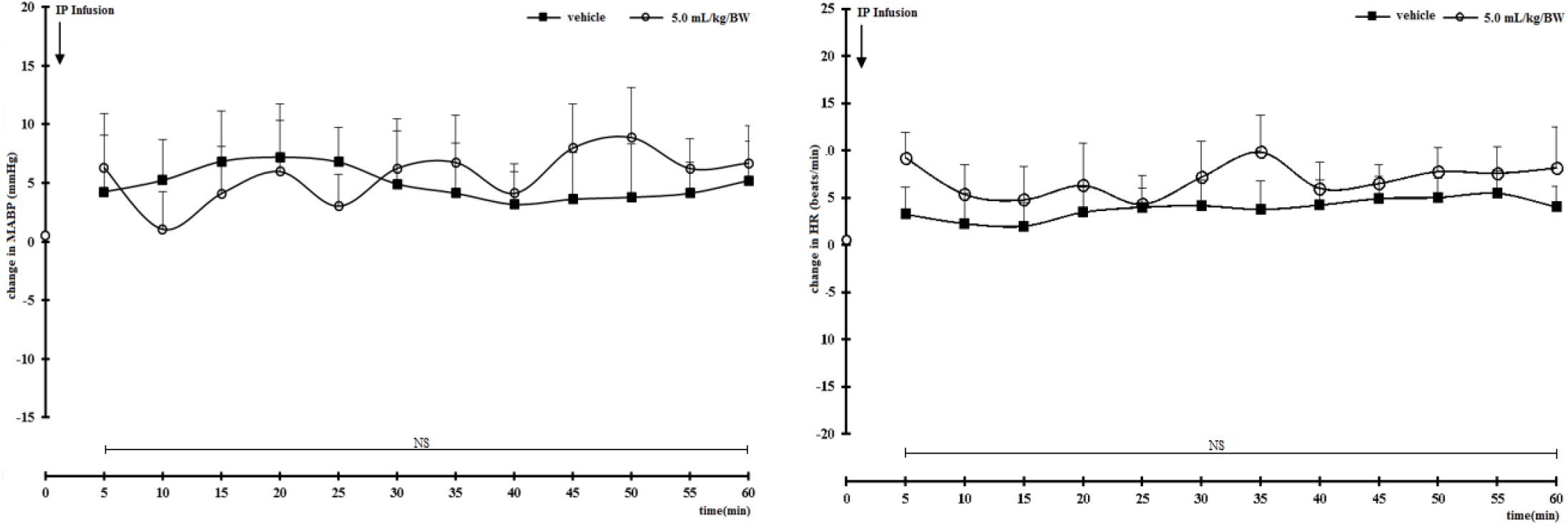
Changes in the mean arterial blood pressure (MABP, mmHg) and heart rate (HR, beats/min) after the intraperitoneal administration of vehicle (saline/ aqueous 0.9 % NaCl n = 5) or methane-rich saline (MRS) (n = 5) at a dose of 5.0 mL/kg/BW in normotensive Sprague Dawley rats. NS – not significant, comparisons between the treatments by ANOVA for repeated measures.

Our findings suggest that methane in contrast to other gasotransmitters such as nitric oxide, hydrogen sulfide or carbon monoxide, does not contribute to arterial blood pressure regulation. However, further studies including chronic experiments in freely moving animals are needed to fully establish possible hemodynamic effects of the gas.

## Materials and Methods

### Animal experiments

The experiments were performed according to Directive 2010/63 EU on the protection of animals used for scientific purposes and approved by the 2nd Local Bioethical Committee in Warsaw (permission number: WAW2/127/2018). Animals were obtained from the Animal Breeding Department of the Medical University of Warsaw and housed in the Central Laboratory of Experimental Animals in group cages with access to standard laboratory chow and water ad libitum. The rats were maintained in a temperature- and humidity-controlled room with a 12/12-hour light– dark cycle. Studies were performed on the male, 18–20-weeks-old, normotensive Sprague Dawley rats (SD). All measurements were performed under general anesthesia with urethane (Sigma-Aldrich) given intraperitoneally (IP) at a dose of 1.5 g/kg of body weight (BW). Before the measurement, rats were implanted with venous and arterial catheters. The arterial catheter was connected to the Biopac MP 150 (Biopac Systems, Goleta, USA) for hemodynamic recordings. The venous catheter was used for the administration of investigated compounds.

### Methane‑Rich Saline Production

With the help of a pressure apparatus, methane was dissolved in saline for 6 h under high pressure (0.4 MPa) to obtain an oversaturated solution. Due to the inevitable de-gassing phenomenon, methane-rich saline MRS was freshly prepared every time before the animal experiments to ensure a steady methane concentration in the injection. The methane concentration in the MRS was determined using gas chromatography (Gas chromatography-9860, Qiyang Co., Shanghai, China).

### Hemodynamic measurements

Measurements started 60 min after the induction of anesthesia and stabilization of hemodynamic parameters. Arterial blood pressure was recorded 20 min at baseline (before treatment) and 60 min after intravenous or intraperitoneal infusions of either vehicle (saline, aqueous 0.9 % NaCl) or methane-rich saline (MRS) at a dose of 0.5 or 1.0 mL/kg BW given by the intravenous (IV) injection and 5.0 mL/kg BW given by the intraperitoneal (IP) injection).

### Data analysis and statistics

The data are expressed as mean+SEM (standard error). The MABP and HR were calculated on the arterial blood pressure tracer using the AcqKnowledge 4.3.1 Biopac software (Biopac Systems, Goleta, USA). For the evaluation of the MABP and HR response within the series, baseline recordings were compared with recordings after administering evaluated compounds using the analysis of variance (ANOVA) for repeated measures. Comparisons between the treatments were performed using the ANOVA. If ANOVA showed a significant difference, a post-hoc Tukey’s test was performed. A value of two-sided p < 0.05 was considered significant. Analyses were conducted using STATISTICA 13.0.

## Author Contributions and Notes

E.Z., D.K., R.O. and M.U. designed research, E.Z., D.K. and M.G.K. performed research, E.Z., D.K. and M.U. analyzed data; and E.Z. and M.U. drafted the manuscript..

The authors declare no conflict of interest.

## Acknowledgments

This study was supported by The National Science Centre of Poland under grant No 2016/22/E/NZ5/00647.

